# Intracellular Delivery of Antibodies for Selective Cell Signaling Interference

**DOI:** 10.1101/2021.10.05.463275

**Authors:** Rebecca L. Hershman, Yamin Li, Feihe Ma, Qioabing Xu, James A. Van Deventer

## Abstract

Many intracellular signaling events remain poorly characterized due to a general lack of tools to interfere with “undruggable” targets. Antibodies have the potential to elucidate intracellular mechanisms via targeted disruption of cell signaling cascades because of their ability to bind to a target with high specificity and affinity. However, due to their size and chemical composition, antibodies cannot innately cross the cell membrane, and thus access to the cytosol with these macromolecules has been limited. Here, we describe strategies for accessing the intracellular space with recombinant antibodies mediated by cationic lipid nanoparticles to selectively disrupt intracellular signaling events. To enable such investigations, we first produced a series of antibody constructs, known as scFv-Fcs, containing additional, genetically encoded negative charges located at the C-termini of the constructs. Preparing proteins with negatively charged motifs has previously been shown to enhance intracellular protein delivery with cationic lipids, but usually for the purpose of genome editing or targeted cell death. We started by generating derivatives of scFv-Fc17, an antibody construct previously reported to bind specifically to signal transducer and activator of transcription 3 (STAT3) phosphorylated at Tyr705 (pYSTAT3). We screened a small number of lipids from our combinatorial lipid library with flow cytometry and found that PBA-Q76-O16B facilitated the most efficient delivery of scFv-Fcs under the conditions tested. In HepG2 cells, we observed up to 60.5% delivery efficacy, while in a STAT3-luciferase reporter cell line up to 71.5% delivery efficacy was observed. These results demonstrated the feasibility of accessing the intracellular space with scFv-Fcs. However, we also note that no more than modest changes were observed upon changing the numbers of negative charges in these constructs during delivery. Characterization of the cytotoxicity, size, and encapsulation efficiency of scFv-Fcs with PBA-Q76-O16B revealed that the constructs were generally well-behaved, with addition of differing quantities of negative charge resulting in at most modest effects. Importantly, functional assays monitoring transcriptional activity in luciferase reporter cell lines and HepG2 cells demonstrated significant reduction of gene expression downstream of pYSTAT3 following delivery of scFv-Fc17 constructs. Together, our results establish the use of recombinantly produced antibodies to selectively interfere with cell signaling events driven by a single posttranslational modification. Efficient intracellular delivery of engineered antibodies opens up possibilities for modulation of previously “undruggable” targets, including for potential therapeutic applications.

## Introduction

Modulating individual protein-protein interactions to investigate how cell signaling cascades impact cell behavior remains a critical challenge in fundamental biology and drug discovery. Numerous therapeutically relevant modalities including nucleic acids, small molecules, peptides, and proteins each provide partial solutions to examining intracellular protein-protein interactions in human cells^1, 2^. Nucleic acid-based strategies (including RNA interference, antisense oligonucleotides, and CRISPR-Cas systems) readily facilitate down- or upregulation of gene expression, but this results in changes to all of the interactions between the protein of interest and its binding partners, not just the targeted interaction. Thus, resulting changes in cell behavior cannot be attributed to individual protein-protein interactions or to signaling events mediated by a particular interaction. Recent advances in small molecule and peptide discovery have shown that these (macro)molecules are capable of selectively disrupting protein-protein interactions^3-6^. While exciting progress is being made to make the discovery of these entities more routine, ensuring that these structures are sufficiently “druglike” remains an ongoing medicinal chemistry challenge^6-8^. In contrast, proteins possess a number of advantages for modulating protein-protein interactions over other modalities. Antibodies and other binding proteins can be engineered to recognize a single target, or even a specific epitope on a target, with subnanomolar affinities and a high degree of specificity^9-11^. This makes them excellent candidates to interact with individual proteins, even down to the level of specifically recognizing posttranslational modifications that drive protein-protein interactions of interest^12^. However, an ongoing challenge in using macromolecules such as antibodies intracellularly is efficiently achieving access to the cytosol^13^. Despite extensive work to devise strategies suitable for intracellular protein delivery in recent years^14^, these efforts must be advanced further in order to routinely utilize proteins to probe intracellular processes.

Delivery vehicles including cell penetrating peptides (CPPs), polymer nanoparticles, and lipid nanoparticles have been explored widely in search of effective intracellular antibody delivery strategies^15, 16^. Each of these vehicles possesses a unique set of advantages and limitations in terms of delivery efficacy, biocompatibility, and impact on antibody properties; these and numerous other considerations have been described extensively in review articles^14, 17, 18^. Further, these strategies commonly require high concentrations of antibody cargo in order to achieve a desired delivery efficiency^19-21^. In early studies, synthetic lipids emerged as promising intracellular delivery agents for antibodies^22-24^. Recent work has also shown that electrostatically assembled complexes formed by mixing cationic lipid-based nanocarriers with negatively charged proteins can efficiently enter the cytosol of human cells while retaining functionality^25^. Investigations by Liu and coworkers showed that fusion of supercharged GFP to genome editing proteins facilitates intracellular delivery mediated by commercially available lipid transfection agents^26^. Liu and coworkers expanded this strategy by screening the human proteome for endogenous, anionic proteins that could enhance intracellular uptake^27^. Studies by Xu and coworkers have further demonstrated that genome editing and cytotoxic proteins modified to be negatively charged can be delivered intracellularly using customized lipid nanoparticles generated through a combinatorial library-based approach^28-31^. However, these promising strategies have not yet been used to deliver antibodies intracellularly to our knowledge. Tsourkas and coworkers recently demonstrated that commercially available lipid transfection reagents support delivery of antibodies with negatively charged polypeptide chains photocrosslinked to antibody Fc domains (Fc: fragment crystallizable)^21^. While these findings are very promising, there remain many outstanding questions about efficient antibody delivery. Two key aspects we seek to investigate are 1) delivering antibody constructs intracellularly without irrelevant fusion proteins or conjugated peptides; and 2) using cationic lipids from combinatorial libraries to enhance antibody delivery beyond what is possible with commercial transfection reagents. These and other outstanding questions highlight opportunities to further enhance intracellular antibody delivery.

In this work, we sought to identify antibody delivery strategies that support specific interference with cell signaling pathways. To accomplish this, we have combined the use of cationic lipid nanoparticles with a series of recombinantly produced antibody constructs known as scFv-Fcs (Figure 1A). The use of recombinant antibodies that were prepared in house allowed us to encode varying levels of negative charge within the constructs to probe potential effects of charge on delivery efficiency and complex formation. A subset of lipids varying in head structure, tail length, and linker composition complexed with scFv-Fcs enabled us to investigate intracellular delivery efficiency. We identified lipids that facilitate efficient scFv-Fc delivery in multiple human cell lines using antibody concentrations as low as 10 nM. Notably, complexes formed with the most promising lipid did not appear to affect cell viability under the conditions tested here. Comparisons of delivery efficiency, cell viability, and particle size as a function of amount of negative charge in the constructs revealed at most modest changes. By preparing scFv-Fcs based on a previously described antibody construct known to interfere with STAT3 signaling (known as scFv-Fc17^32^), we were able to evaluate whether delivered constructs could disrupt phosphorylation-mediated STAT3 dimerization in the JAK-STAT pathway (Figure 1B). Interfering with this pathway by blocking dimerization is well-suited for an antibody-based approach, but considerably more challenging for strategies using small molecules or peptides and essentially impossible with nucleic acid-based knockdown or knockout strategies. Luciferase assays conducted in a series of reporter cell lines resulted in specific interference with STAT3 signaling following delivery of scFv-Fc17 constructs, but no changes in signaling were detected following delivery of control constructs or in cell lines that report on signaling activity via STAT1 or NFκB pathways. Finally, we used RT-qPCR in the HepG2 cell line to further validate the ability of delivered scFv-Fcs to interfere with signaling via the STAT3 pathway. These experiments revealed clear changes in gene expression levels following delivery of scFv-Fc17 constructs but no changes following delivery of control constructs. Taken as a whole, our data provides strong evidence for the efficient delivery of recombinant antibody constructs and specific interference with an intracellular signaling pathway. Routine access to the intracellular space with engineered antibodies and other binding proteins will facilitate new opportunities for probing the roles of individual protein-protein interactions and posttranslational modifications. These molecular-level investigations are invaluable for deepening our understanding of basic cellular processes and for identifying potential therapeutic strategies.

**Figure 1.**
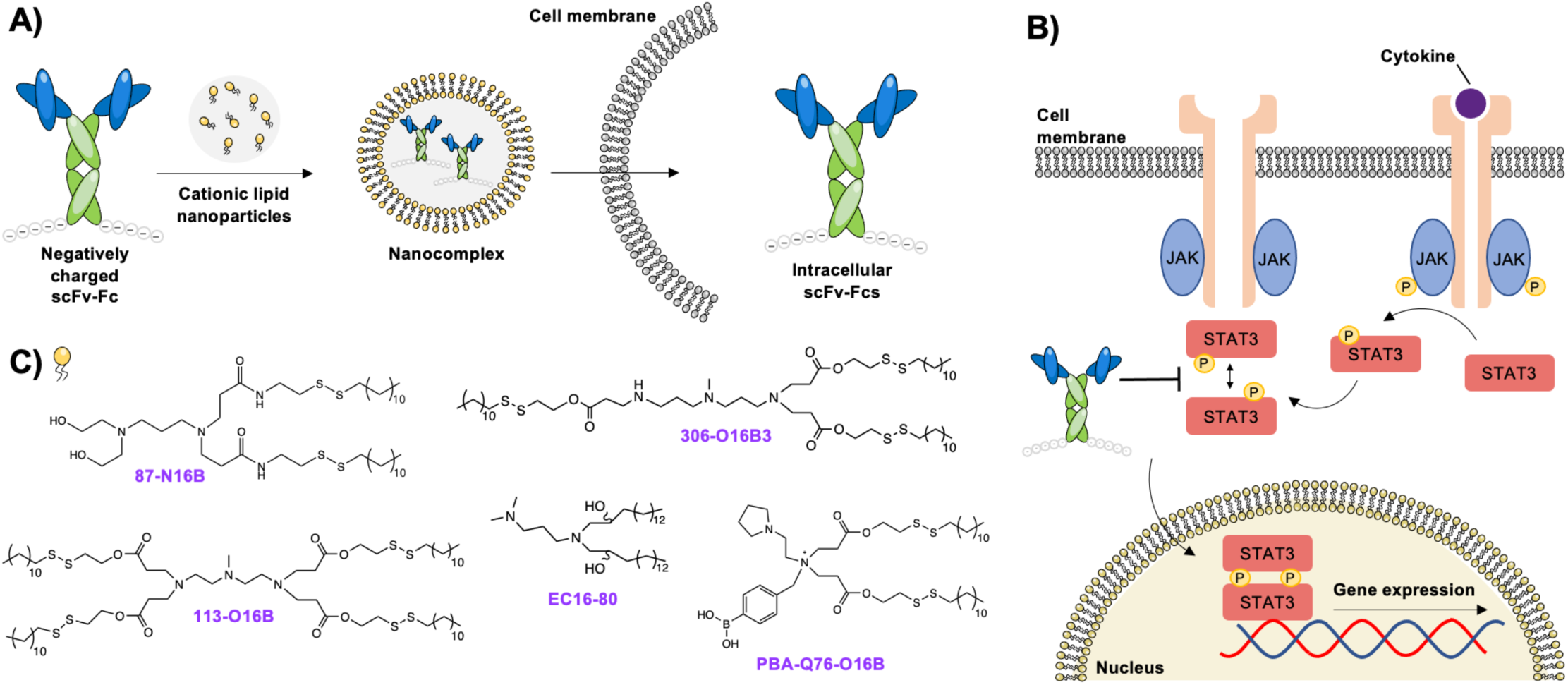
Overview of delivery strategy and targeted intracellular signaling pathway. A) ScFv-Fc antibody constructs are complexed with cationic lipids to form submicrometer particles. Treatment of mammalian cells with these particles leads to intracellular access and release of scFv-Fc. B) Summary of signaling via JAK/STAT pathway. Specific interference with signaling can be achieved using the antibody construct scFv-Fc17, which specifically binds to pYSTAT3 and disrupts dimerization. C) Chemical structures of lipids used in this study.

## Results and Discussion

### ScFv-Fc preparation and validation

To evaluate strategies for leveraging intracellular antibody delivery to disrupt intracellular processes, we sought to identify an antibody construct with known ability to interfere with a signaling pathway. We found the previously described construct named scFv17, which specifically binds to STAT3 phosphorylated at Tyr 705 (pYSTAT3)^32^. Rhee and coworkers demonstrated that transient transfection of an scFv-Fc form of scFv17 in HepG2 cells led to disruption of signaling downstream of pYSTAT3 in the JAK/STAT pathway (see Figure 1B for schematic of pathway^33, 34^). In order to use scFv17 in this work, we purchased a yeast codon-optimized form of the DNA encoding the antibody fragment and cloned it into the secretion vector pCHA-FcSup-TAG^35^. This resulted in pCHA-FcSup-scFv17-TAG, a yeast secretion plasmid construct that encodes for scFv-Fc17, where scFv17 is expressed as an N-terminal fusion to an IgG1 fragment crystallizable (Fc) domain. We also utilized a control scFv-Fc secretion construct where the scFv targeting pYSTAT3 was replaced with a binder to fibroblast activation protein (FAP; scFv-Fc construct termed FAP-Fc), a membrane protein that is not expected to be present in the cytoplasm of human cells (see *Supplementary Methods* for details). In addition, we also prepared versions of the scFv-Fcs in which 5 or 15 additional glutamic acid residues were encoded at the C-terminus of the Fc, allowing for initial investigations into the role of negative charge in delivery of these recombinant constructs (SI Figure 1). ScFv-Fcs containing 5 additional glutamic acid residues are referred to as scFv-Fc17-5E and FAP-Fc-5E, while constructs containing 15 additional glutamic acid residues are referred to as scFv-Fc17-15E and FAP-Fc-15E. Each of these scFv-Fcs were secreted and purified from yeast culture as described in *Supplementary Methods* (SI Figure 1).

To confirm the binding activities of the purified constructs, we conducted Western blotting experiments using HepG2 cell lysates and STAT3-luciferase cell lysates using scFv-Fc17 and FAP-Fc protein constructs as primary antibodies. These experiments utilized lysates derived from cultures treated with cytokines known to stimulate STAT3 signaling as well as lysates derived from unstimulated controls. Bands were detected at the expected molecular weight of pYSTAT3 (∼88 kDa) for both cell lines exclusively following cytokine stimulation and when scFv-Fc17 constructs were used as primary antibodies. This is consistent with the specific recognition of the STAT3 isoform observed with a commercially available anti-pYSTAT3 antibody (SI Figure 2, 3). Further, no construct binding was observed when the control FAP-Fc constructs were used to probe the membranes. Moreover, as expected, the addition of 5 or 15 glutamates to scFv-Fc17 did not affect the recognition of pYSTAT3 or increase nonspecific binding for any construct. Overall, these results confirm that constructs based on scFv-Fc17 and FAP-Fc appear to be well-behaved, with scFv-Fc17 constructs specifically binding to pYSTAT3 and control constructs exhibiting no detectable binding to cell lysates.

### Identifying preferred delivery conditions with FITC-labelled scFv-Fcs

To begin investigating lipid-based delivery of scFv-Fcs, we complexed fluorescently-labelled antibody constructs with selected lipids from our combinatorial library (Figure 1C)^28, 29, 36, 37^ and used flow cytometry to evaluate scFv-Fc uptake in HepG2 cells. In previous work, these candidate lipids showed favorable intracellular delivery properties for genome editing proteins, cytotoxic proteins, and nucleic acid-based cargo, and thus we sought to determine if these efficient delivery properties could be extended to antibody constructs. We used the HepG2 cell line for the initial screen because the STAT3 signaling pathway can be stimulated in these cells, with prior work showing that specific signaling via pYSTAT3 can be assayed^32, 38^. For an initial evaluation of delivery, scFv-Fc17, scFv-Fc17-5E, and scFv-Fc17-15E were labelled with amine-reactive fluorescein isocyanate (FITC) and complexed with lipids in a 1:1 mass ratio. The resulting nanocomplexes were added to tissue culture wells containing HepG2 cells at a final concentration of 35 nM antibody construct. Following 24 hours of incubation, cells were analyzed using flow cytometry, and the percentage of FITC-positive cells for each condition was determined (Figure 2). Under these conditions, the highest percentages of FITC-positive cells were observed when using lipids 306-O16B3^36^ and PBA-Q76-O16B^37^. In some cases, we observed modest increases in the percentage of FITC-positive cells as negative charge in the construct was increased. These small, or in some cases negligible, effects are consistent with the recent findings of Tsourkas and coworkers, where they reported that substantial improvements in antibody delivery required at least 20 negative charges to be linked to native antibodies^21^. Overall, these preliminary findings indicated that lipids 306-O16B3 and PBA-Q76-O16B mediate efficient antibody delivery under the conditions we tested, motivating us to investigate additional experimental parameters in search of enhanced cellular uptake.

**Figure 2.**
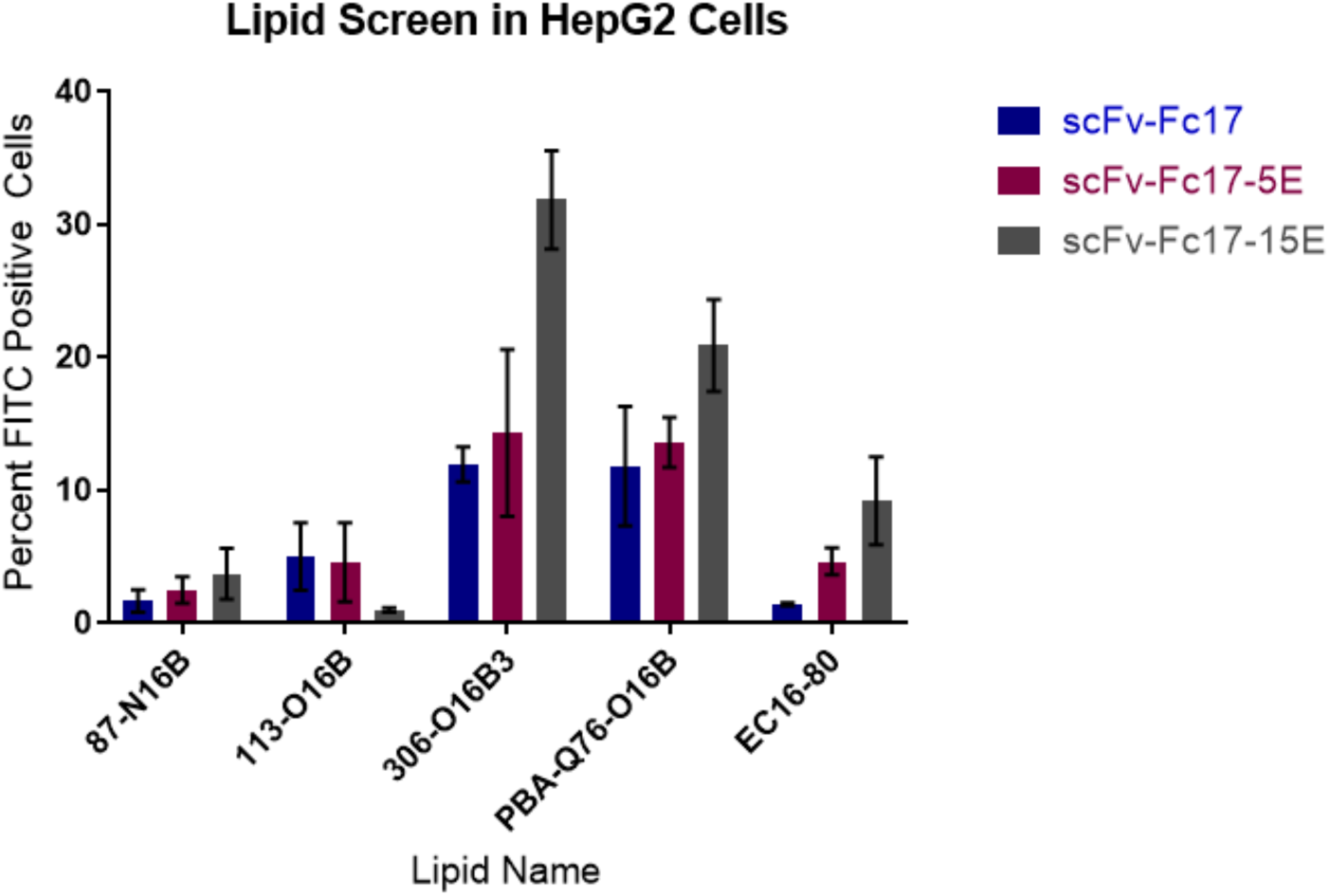
Initial lipid screen with HepG2 cells assessing scFv-Fc delivery with 5 lipids from the combinatorial lipid library. HepG2 cells were incubated for 24 hours with nanocomplexes formed in a 1:1 mass ratio of scFv-Fcs to lipid with the protein dose constant at 35 nM in media. Following incubation, cells were washed three times with 1× PBS containing 20 U/mL heparin and then analyzed via flow cytometry for detection of FITC-labelled scFv-Fcs. Data shown is a mean of three replicates (n = 3) where error is the standard deviation from the mean.

Next, we varied protein to lipid mass ratios during preparation of nanocomplexes in search of more efficient delivery conditions. Previous work has shown that adjusting the protein to lipid ratio can be an effective method to enhance intracellular delivery^39^. In order to evaluate high lipid mass-to-protein complexation conditions, we lowered the final protein concentration to 10 nM during incubation with cells. This lower protein concentration allowed us to evaluate higher lipid loading while reducing the risk of cellular toxicity. Keeping the protein dose constant at 10 nM, scFv-Fc17 and negatively charged constructs were combined with various masses of lipid at the indicated ratios using the lipids 306-O16B3 and PBA-Q76-O16B (Figure 3, SI Figure 3). These experiments were conducted using both HepG2 cells and STAT3-luciferase cells to determine preferred delivery conditions for both cell lines. Delivery following complexation with lipid PBA-Q76-O16B resulted in increasing percentages of FITC-positive cells as increasing lipid masses were used in both HepG2 and STAT3-luciferase cell lines (Figure 3). When the protein to lipid mass ratio reached 1:8, both cell lines showed high percentages of FITC-positive cells for all three scFv-Fc17 constructs investigated here. For HepG2 cells, 48.9% (±12.1), 58.4% (±8.6), and 60.5% (±20.5) FITC-positive cells were observed for 0, 5, and 15 additional negative charges, respectively. With STAT3-luciferase cells, we observed 70.8% (±2.7), 71.5% (±7.3), and 71.2% (±2.4) FITC-positive cells for the respective constructs. In contrast to these findings, increases in lipid mass using lipid 306-O16B3 did not result in consistently higher percentages of FITC-positive cells in either cell line tested (SI Figure 3). For both lipids systematically evaluated here, delivery with 10 nM antibody construct resulted in modest or undetectable changes as negative charge in the constructs was increased, consistent with our initial delivery tests at higher concentrations. Irrespective of charge, the data clearly indicates that the highest delivery efficiency is observed in both cell lines with 10 nM antibody concentration complexed with the lipid PBA-Q76-O16B in a 1:8 mass ratio of protein to lipid.

**Figure 3.**
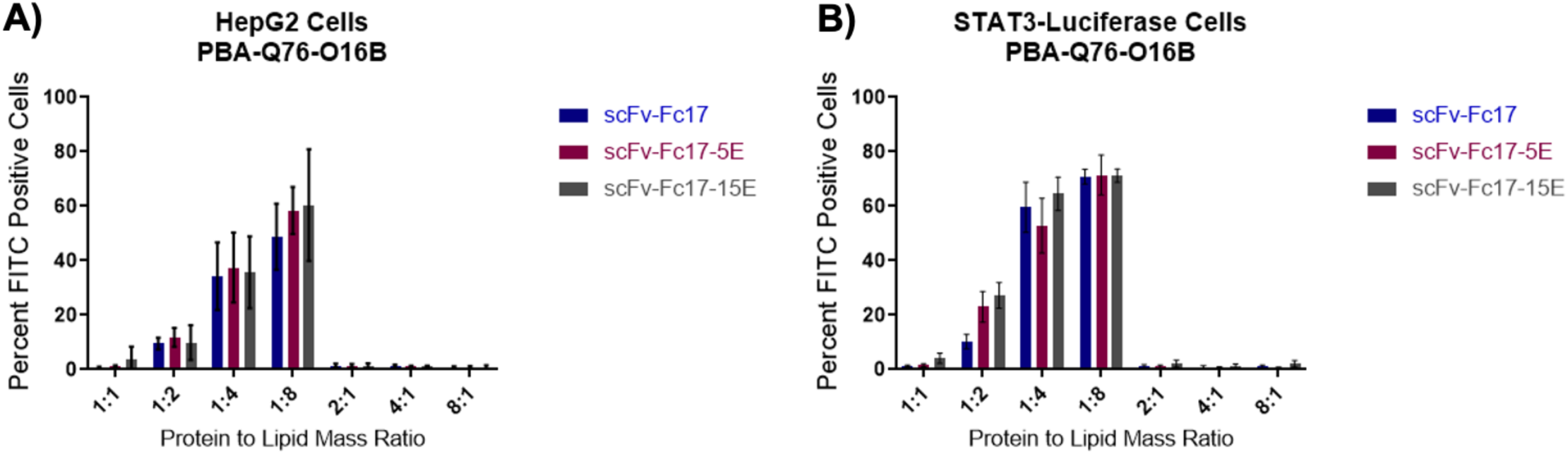
Flow cytometry analysis of changes in protein to lipid mass ratio for delivery. A) HepG2 cells and B) STAT3-luciferase cells were incubated with 10 nM scFv-Fcs complexed at indicated mass ratios with lipid PBA-Q76-O16B for 24 hours. Following incubation, cells were washed three times with 1× PBS containing 20 U/mL heparin and then analyzed via flow cytometry for detection of FITC-labelled scFv-Fcs. Data shown is a mean of two independent experiments each containing three technical replicate wells where error is the standard deviation from the mean (n = 6).

In order to ensure that we could conduct well-controlled investigations of cell signaling, we further investigated whether the preferred delivery conditions described above result in efficient delivery with control luciferase reporter cell lines and FAP-Fc control proteins. Both STAT1-luciferase and NFκB-luciferase reporter lines provide opportunities for assessing if delivered proteins interfere specifically with the STAT3 pathway (see *Supplementary Methods* for details on cell lines). Flow cytometry experiments with the STAT1-luciferase cell line and scFv-Fc17 constructs complexed with PBA-Q76-O16B resulted in similar levels of FITC-positive cells in comparison to the STAT3-luciferase line (SI Figure 4a). We also made attempts to evaluate the delivery properties of the NFκB-luciferase cell line, but this line does not possess adequate adherence to tissue culture surfaces to survive the stringent washes necessary for analysis with flow cytometry. As such, given the similar delivery efficiencies observed with the closely related STAT3- and STAT1-luciferase lines, we still used the NFκB-luciferase cell line in subsequent experiments.

The FAP-Fc series of control proteins contain the distinct anti-FAP scFv but identical Fc regions to those found in the scFv-Fc17 series of constructs. These control constructs were complexed under the preferred conditions and delivered to HepG2 cells, STAT3-luciferase cells, and STAT1-luciferase cells to evaluate construct delivery. SI Figure 4b demonstrates that these constructs are delivered intracellularly with similar efficiency to their scFv-Fc17 counterparts under the same conditions for each cell line. Taken as a whole, our flow cytometry analyses of scFv-Fc delivery demonstrates that our system is well-behaved across multiple constructs and cell lines. Although we were not able to observe robust changes in delivery efficiency using additional negative charge, we were able to identify conditions under which we observed consistent intracellular access using all six scFv-Fc constructs in multiple cell lines. We have all of the necessary controls to comprehensively evaluate whether the delivery conditions we have identified will facilitate interference with STAT3-mediated signaling.

### Characterization of protein-lipid nanocomplexes

Following the determination of conditions that support high-level antibody construct delivery, we moved on to evaluate the cytotoxicity, size, and encapsulation efficiency of protein-lipid nanocomplexes under these conditions. To measure the cytotoxicity of the nanocomplexes, we treated each cell line with scFv-Fc nanocomplexes assembled in a 1:8 mass ratio of protein to PBA-Q76-O16B, as well as some samples with PBA-Q76-O16B alone (Figure 4A). We found that almost all samples maintained viabilities above 85% following incubation for 24 hours, suggesting that the delivery conditions we identified result in no overt toxicities on the cell lines tested here.

**Figure 4.**
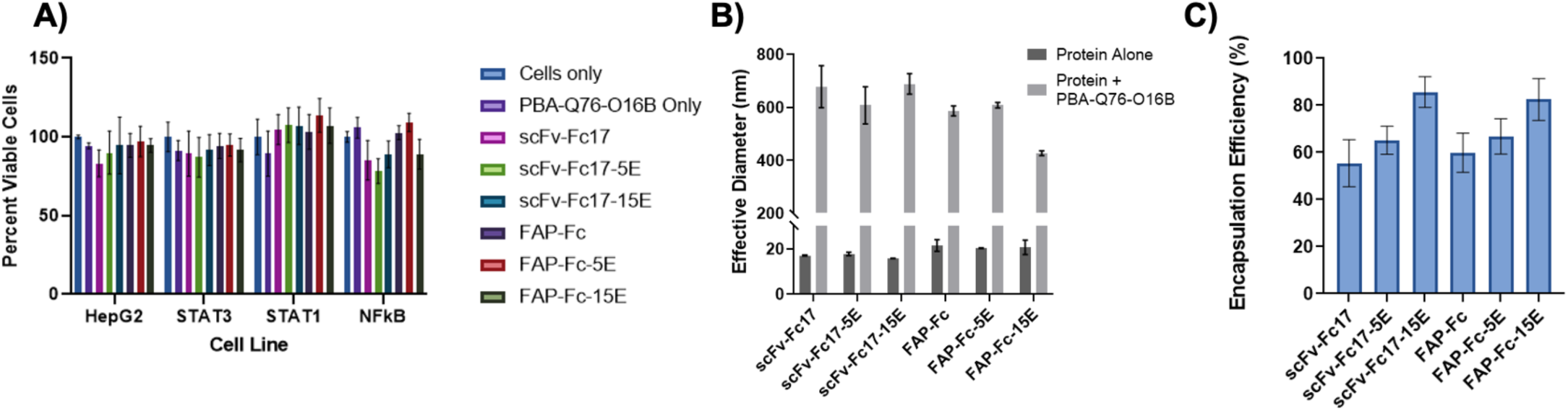
Nanoparticle characterizations. A) Cell viability assays for all proteins used in this study. Cells were treated for 24 hours with PBA-Q76-O16B alone or with indicated scFv-Fc complexed in a 1:8 mass ratio of protein to PBA-Q76-O16B at a 10 nM final protein concentration. Cell viability was assessed using CellTiter-Fluor Cell Viability Assay. Data shown is a mean of individual samples (n = 3) and error is represented by the standard deviation propagated through analysis. B) Dynamic light scattering measurements evaluating effective diameter of proteins alone as well as proteins complexed in a 1:8 protein to lipid mass ratio with PBA-Q76-O16B. Data shown is a mean of individual samples (n = 3) and error is represented by the standard deviation. C) Encapsulation efficiency of proteins complexed with PBA-Q76-O16B. Nanocomplexes were formed by mixing PBA-Q76-O16B in a 1:8 protein to lipid mass ratio with indicated scFv-Fcs in 1× PBS. After 15 minutes, nanoparticles were spun down at 21,000 rcf for 45 minutes and the protein concentration in the supernatant was measured by BCA Assay. The percent of protein encapsulated was determined by mass balance. Data shown is a mean of three independent replicates (n = 3) and error is represented as the standard deviation propagated through analysis.

We also evaluated the size and protein encapsulation efficiencies obtained upon complexing PBA-Q76-O16B with all six antibody constructs evaluated in this work. Dynamic light scattering (DLS) measurements were used to estimate the effective sizes of the complexes (Figure 4B). For the size measurements, scFv-Fc nanocomplexes were assembled in a 1:8 mass ratio of protein to PBA-Q76-O16B. The lipid alone in solution was determined to have a particle size of approximately 100 nm, which is consistent with other reports using this lipid^37^. All measurements of protein-lipid complexes fell within the range of 400 to 700 nm, with no clear correlations between the measured size and the number of negative charges present in the protein constructs. In contrast to characterizations of protein size, measurements of encapsulation efficiency revealed that the percentage of protein contained in the nanocomplexes increases as the charge of the protein construct increases (Figure 4C). scFv-Fc17 and FAP-Fc were encapsulated at percentages of 55.5% (±10.0) and 57.6% (±8.3), respectively, while scFv-Fc17-5E and FAP-Fc-5E were encapsulated 65.2% (±5.9) and 68.8% (±7.5). ScFv-Fc17-15E and FAP-Fc-15E were encapsulated at 85.6% (±6.5) and 82.5% (±8.8), respectively. Overall, these data indicate that cells retain viability following treatment with these nanocomplexes, the complexes are hundreds of nanometers in effective diameter, and encapsulation efficiencies increase with increasing charge in protein construct.

### Signaling interference with scFv-Fc17 constructs in luciferase reporter cell lines

Having identified efficient delivery conditions that preserve cell viability, we initiated studies to evaluate changes in signaling via the STAT3 pathway following delivery of scFv-Fc17 constructs. Because encapsulation efficiencies appeared to be higher for the 5E and 15E scFv-Fcs, we only used these constructs during signaling assays. To directly investigate interference with STAT3 signaling via pYSTAT3, we utilized a STAT3-luciferase line that expresses firefly luciferase in response to stimulation with the cytokine oncostatin M (OSM; see *Supplementary Methods* for details). For delivery experiments with this line, we expect to see reduced luciferase activity following delivery with scFv-Fc17 constructs. Using the preferred conditions identified above, incubation of STAT3-luciferase cells with PBA-Q76-O16B complexed with scFv-Fc17-5E or scFv-Fc17-15E for 24 hours prior to stimulation with OSM led to luciferase activity levels at 59.6% (±14.9) and 59.8% (±15.0) of stimulated control cultures (Figure 5a). In contrast, incubation and stimulation under the same conditions using FAP-Fc-5E and FAP-Fc-15E resulted in no significant change in luciferase activity in comparison to an OSM-only treated control (Figure 5a). Together, these data demonstrate that, even at an effective concentration of 10 nM, complexed scFv-Fc17 constructs decrease STAT3-dependent luciferase activity in a reporter cell line, while irrelevant FAP-Fc constructs do not.

**Figure 5.**
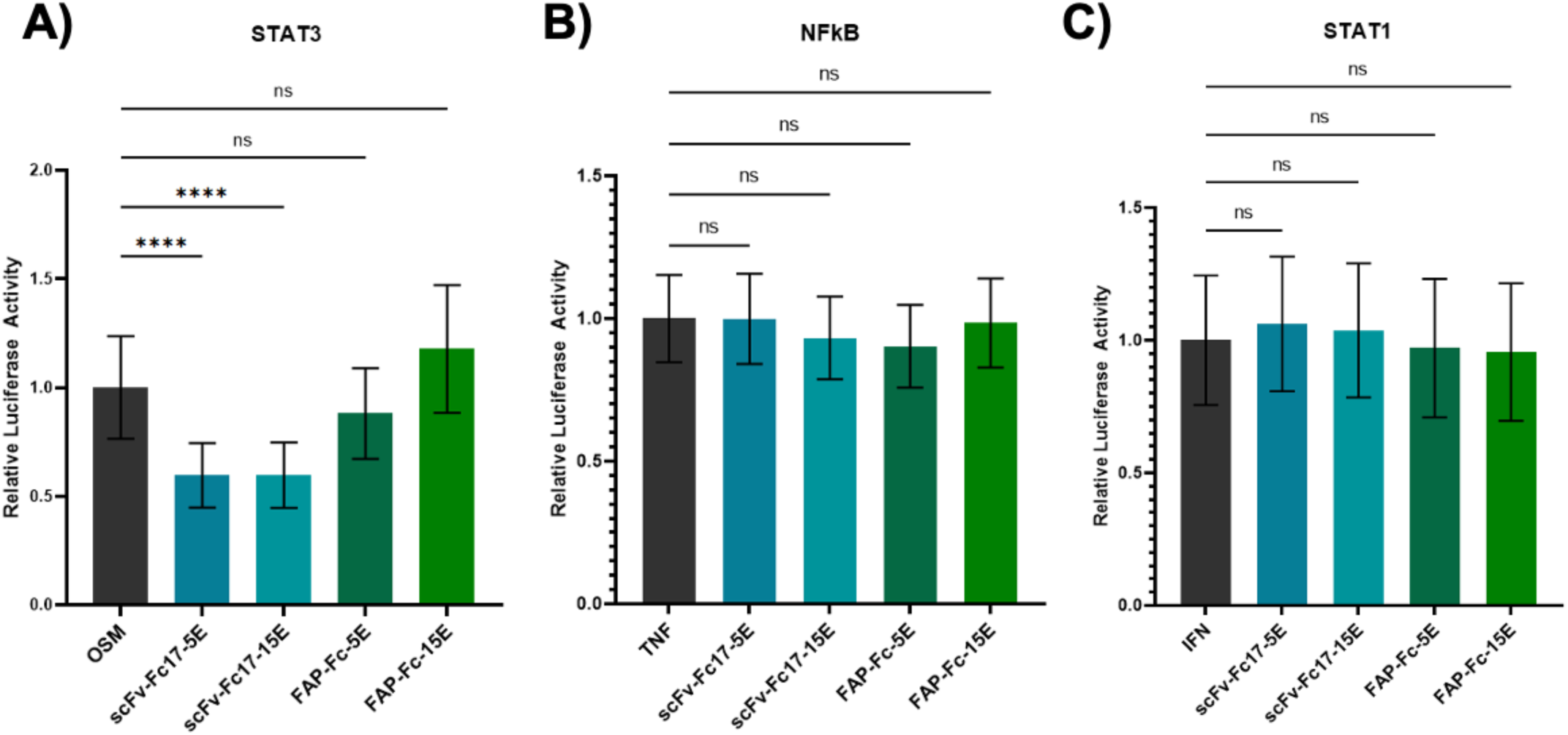
Luciferase assay results. A) STAT3-luciferase cells were incubated with indicated scFv-Fcs at a 10 nM final concentration complexed with lipid PBA-Q76-O16B in a 1:8 mass ratio for 24 hours in media. Cells were then stimulated with 10 ng/mL OSM for 5 hours to activate the luciferase pathway. BrightGlo Luciferase Assay System was used to quantify luciferase expression, and luminescence was detected on a SpectraMax i3x microplate reader. Luminescent data was normalized to viable cell counts using the CellTiter-Fluor Cell Viability Assay. B) Same as in A) but for NFκB-luciferase cells stimulated with 10 ng/mL TNFα. C) Same as in A) but for STAT1-luciferase cells stimulated with 10 ng/mL IFNγ. Data shown is a mean of individual samples (n = 14) normalized to the stimulated-only control, and error is represented by the standard deviation propagated through analysis. Statistical analysis is from a one-way ANOVA where **** indicates p ≤ 0.0001 and ns indicates no significance.

To verify that the changes in signaling we detected could be attributed specifically to the disruption of signaling via STAT3, we conducted additional experiments using STAT1-luciferase cells (derived from the same parental line as for the STAT3-luciferase line) and NFκB-luciferase cells (derived from 293 cells)^40^. Signaling in the STAT1-luciferase and NFκB-luciferase lines can be achieved using interferon-γ (IFNg), and tumor necrosis factor-α (TNFa), respectively, via the signaling pathway for which the cell line is named. For complexed scFv-Fc17-5E, scFv-Fc17-15E, FAP-Fc-5E, and FAP-Fc-15E, treatment of STAT1-luciferase and NFκB-luciferase lines prior to cytokine stimulation resulted in insignificant changes in signaling via these pathways (Figure 5B, C). In the case of STAT1, this lack of signaling is especially noteworthy because STAT1 and STAT3 proteins are known to heterodimerize^41^. The absence of changes observed here indicates that delivery of constructs that bind to and interfere with signaling via pYSTAT3 exhibit no overt off-target interference with closely related pathways. Taken together, these luciferase assays demonstrate that our delivery strategy facilitates interference with an intracellular signaling pathway controlled by highly specific phosphorylation events.

### RT-qPCR analysis of genes downstream pYSTAT3 in HepG2 cells

After observing specific reduction in STAT3-dependent signaling using reporter cell lines, we attempted to use scFv-Fc delivery in HepG2 cells to interfere with gene expression downstream of pYSTAT3. To provide a stringent test of our delivery strategy, all experiments were performed following treatment of cells with the modest 10 nM effective concentration of scFv-Fcs in complex with PBA-Q76-O16B as described above. Using RT-qPCR, we monitored expression levels of three genes: haptoglobin, serum amyloid A 1 (SAA1), and JunB. These genes are known to be expressed at higher levels in HepG2 cells following stimulation of the JAK/STAT3 pathway with interleukin-6 (IL-6)^42-45^. In the experiments conducted here, HepG2 cells were incubated for 24 hours with complexed antibody constructs, washed once with 1× PBS, and then stimulated with 50 ng/mL IL-6 in media for 90 minutes. Following IL-6 treatment, total RNA was extracted for RT-qPCR analysis (see *Supplementary Methods* for details). The expression levels of the three genes differed from one another as expected (Figure 6). In the case of SAA1, we observed the highest fold increase of gene expression following IL-6 stimulation (Figure 6A), and significant reduction in expression levels following treatment with scFv-Fc17-5E and scFv-Fc17-15E. Under the conditions investigated here, expression of SAA1 was decreased 2.43- and 2.53-fold, respectively. In contrast, incubation of HepG2 cells with FAP-Fc-5E and FAP-Fc-15E did not show significant changes in SAA1 expression. Taken as a whole, these data show that delivery with scFv-Fc17 constructs reduced SAA1 expression following IL-6 stimulation, while delivery with FAP constructs did not.

**Figure 6.**
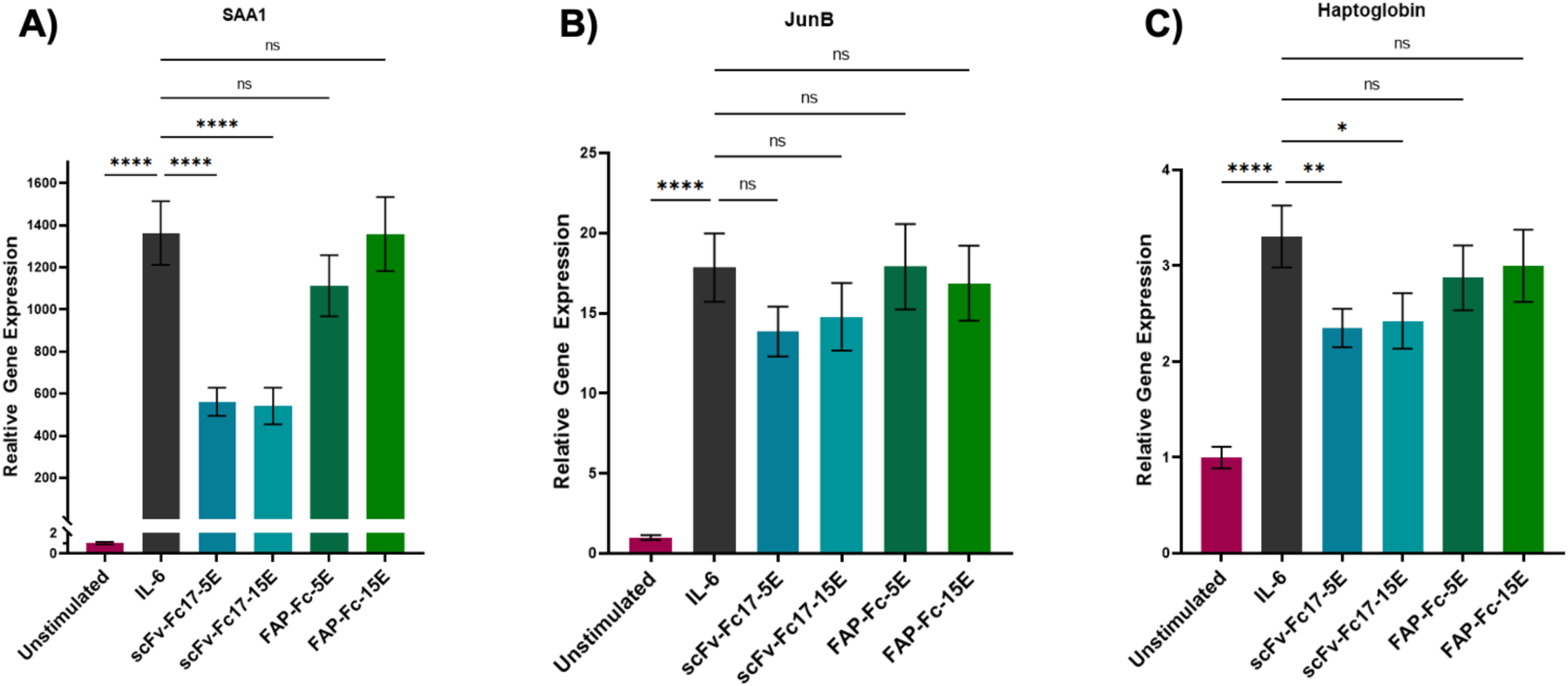
RT-qPCR assay results. Analysis for A) SAA1, B) JunB, and C) haptoglobin. HepG2 cells were incubated with indicated scFv-Fcs at a 10 nM final concentration complexed with lipid PBA-Q76-O16B in a 1:8 protein to lipid mass ratio for 24 hours in media. Cells were stimulated with 50 ng/mL IL-6 for 90 minutes to activate pYSTAT3 pathway. Total RNA was extracted with TriZol, reverse transcribed with SSIV First Strand Synthesis System, and amplified using PowerUp SYBR Green Master Mix with primers listed in Supplementary Information. qPCR data was analyzed using the ΔΔC_T_ method using HPRT as a housekeeping gene. Data shown is the relative gene expression normalized to the unstimulated control, and error is represented by the standard deviation propagated through analysis (n = 3). Statistical analysis is from a one-way ANOVA where **** shows p ≤ 0.0001, ** shows p ≤ 0.01, * shows p ≤ 0.05, and ns shows no significance.

We performed similar evaluations of changes in JunB and haptoglobin expression levels following treatment of cells with antibody constructs. These genes exhibit one to two orders of magnitude changes in expression levels following IL-6 stimulation, in contrast to the three orders of magnitude observed with SAA1. Under the delivery conditions used here, we observed small decreases in JunB expression following treatment with scFv-Fc17-5E (1.29-fold) and scFv-Fc17-15E (1.21-fold) (Figure 6B). Further, treatments with FAP-Fc-5E and FAP-Fc-15E resulted in 0.98-fold and 1.06-fold decreases, respectively. However, we note that these modest changes in JunB expression were not determined to be statistically significant using a one-way analysis of variance (ANOVA). One possible reason for the lack of clear statistical differences in this case could be the small changes in JunB expression levels following stimulation, potentially making it more challenging to observe significant changes in gene expression. On the other hand, for haptoglobin we were able to observe statistically significant decreases in gene expression following delivery with both scFv-Fc17-5E and scFv-Fc17-15E, even though the expression level changes by only several-fold following stimulation (Figure 6C). Treatment of HepG2 cells with complexed scFv-Fc17 constructs prior to stimulation resulted 1.41- and 1.36-fold decreases, respectively. Statistical analysis using a one-way ANOVA indicated that in this experiment the gene expression decreases significantly compared to the control. Incubation with complexed FAP-Fc constructs prior to stimulation resulted in 1.15- and 1.10-fold decreases with the 5E and 15E proteins, respectively; these changes were not determined to be statistically significant. Because the changes in expression levels we observed were relatively small, we performed additional statistical analysis on these data following a different normalization procedure (Figure SI 6), and repeated these full experiments on a different day (Figure SI 7, 8). Across the two experiments and analysis methods, we consistently observed large, statistically significant changes in SAA1 expression levels following treatment with scFv-Fc17 constructs. Statistically significant changes in JunB and haptoglobin expression levels following delivery with scFv-Fc17 constructs were observed in several, though not all, combinations of experiments and analyses reported here. Ultimately, these results show that intracellularly delivered scFv-Fc17 constructs, at a final concentration of only 10 nM, were able to inhibit pYSTAT3 and reduce expression of downstream genes.

## Conclusions

In this work, we were able to demonstrate the utility of our intracellular delivery platform using a combination of recombinantly produced scFv-Fcs engineered to contain additional negative charges and cationic lipid nanoparticles. Using flow cytometry-based experiments, we identified preferred conditions for delivery and characterized the nanocomplexes formed under these conditions. Evaluation of the size of each nanocomplex using dynamic light scattering did not reveal any trends based on the various charges of different scFv-Fcs, while both flow cytometry data and encapsulation efficiency experiments revealed modest changes in both delivery efficiency and encapsulation percentage. In both cases, an increase in negative charge led to small increases in the percentage of protein encapsulated by the lipids as well as delivery efficiency. With inhibitory assays using scFv-Fc17 constructs, we saw significant decreases in gene expression downstream of pYSTAT3 in both STAT3-luciferase cells as well as HepG2 cells. Overall, these experiments represent an important proof-of-concept showing effective delivery of inhibitory scFv-Fcs to regulate a signaling pathway.

Using antibody concentrations as low as 10 nM, we observed robust signaling interference across multiple cell lines following scFv-Fc delivery. In potential therapeutic applications, low antibody concentrations are highly favorable since they reduce the possibility of systemic toxicity. Although we did not observe complete shutdown of the pYSTAT3 signaling pathway under the conditions that we tested, further exploration can be done with this system to determine which experimental parameters would enable greater reductions in signaling. Taken together, our findings provide opportunities to better understand intracellular signaling pathways using antibodies or other specific binding proteins. The functions of many protein-protein interactions remain unknown in signaling cascades, and the ability to probe them with intracellularly delivered, highly specific protein constructs enables controlled explorations. With the exquisite specificities of binding proteins such as scFv17, such investigations are capable of reaching the level of individual posttranslational modifications involved in protein-protein interactions known to drive signaling processes^12, 46^. The platform we have established for intracellular delivery of antibody constructs will enable further exploration of intracellular signaling pathways to better understand molecular mechanisms and potentially elucidate disease targets.

A major unsolved challenge for intracellular protein delivery is accommodating the variable charge and hydrophobicity of proteins, which make it challenging to create a universal delivery method. The delivery strategy we describe in this work involves a combination of several technologies including recombinant antibody production, genetic engineering, and custom lipid synthesis that together have the potential to enable generalization to a wide variety of intracellular targets. Firstly, the use of lipids generated from a combinatorial library enables screening of thousands of possible candidates for desirable delivery properties. Xu and coworkers have demonstrated extensive use of these lipids for a variety of delivery cargo in a variety of cell types *in vitro* as well as *in vivo*. Secondly, recombinant protein production, here scFv-Fc production in yeast, enables us to use standard cloning techniques to generate scFv-Fcs that are specific to desired intracellular targets, provided that the binding properties of the protein have been well characterized. Further, the use of recombinant production strategies allowed us to genetically encode negative charges in our antibody constructs to investigate how such charges may alter delivery properties and signaling interference. Though the studies reported here did not reveal a strong enhancement of delivery with additional negative charges, future work in this area could involve changing the amount or location of negative charge to further investigate strategies to improve delivery efficiencies and thus enhance signaling interference. Recently, our lab demonstrated that we can secrete scFv-Fcs from yeast that contain noncanonical amino acids (ncAAs) at site-specific locations throughout the construct^47^. Some of these ncAAs contain functional groups that are amenable to “click” chemistry reactions^47-49^ that we can leverage to add negatively charged functionalities at selected positions throughout the construct. The constructs in the present work contain additional negative charges only at the C-terminus, but as far as we are aware, no investigations have been made into whether the introduction of negative charge at different locations impact delivery efficiency. Together, these complementary technologies will allow us to investigate conditions under which we can routinely access the intracellular space with antibodies. Combining custom lipid synthesis technology and protein engineering provide important opportunities to modulate the functions of intracellular targets that have previously been considered “undruggable.”

## Acknowledgements

We would like to thank the Frank Lab at Dana Farber Cancer Institute for providing us with the three luciferase reporter cell lines used in this study (STAT3-, STAT1, and NFκB-luciferase). This research was supported by a grant from the National Science Foundation (DMR-1807415, BMAT), and by Tufts University startup funds (to J.A.V.). The content of this work is solely the responsibility of the authors and does not necessarily represent the official views of the National Science Foundation or Tufts University.

## Supplementary materials

Additional experimental details, materials, and figures are provided in the Supporting Information file.

