## Supplementary Information for "Intracellular Delivery of Antibodies for Selective Cell Signaling Interference"

### Supplementary Methods

scFv-Fc Plasmid Construction

In order to produce soluble scFv-Fcs, a previously reported yeast secretion plasmid was used: pCHA-FcSup-TAG[1]. All scFv sequences are listed in SI Table 1, and all primer sequences for cloning are listed in SI Table 2. PCRs were performed with Q5 high fidelity polymerase (New England Biolabs), and constructs were sequence verified via Sanger sequencing (Quintara Biosciences).

To generate pCHA-FcSup-scFv17-TAG, which encodes for scFv-Fc17, we took the scFv17 amino acid sequence reported by Koo et al.[2] and optimized the codons for translation in yeast. The optimized gene fragment was synthesized by Integrated DNA Technologies (IDT), and we amplified it using primers scFv17-FP and scFv17-RP. We then used Gibson Assembly to combine the amplified fragment with pCHA-FcSup-TAG digested with NheI and XmaI.

pCHA-FcSup-FAP0.5b.13-TAG, which encodes for FAP-Fc, was previously reported[1].

pCHA-FcSup-scFv17-5E-TAA, which encodes for scFv-Fc17-5E, was constructed by first modifying the portion of pCHA-FcSup-scFv17-TAG that encodes the C-terminus of the construct to contain 5 additional glutamic acid codons with a modified TAA codon for termination. We started by PCR amplifying two fragments from pCHA-FcSup-scFv17-TAG. One fragment was generated by using primers scFv17-FP and 5E-scFvFc17-R1, and the other was generated by using primers 5E-scFvFc17-F2 and 5E-scFvFc17-R2. These two segments were recombined with the pCHA-FcSup-scFv17-TAG vector digested with NheI and XhoI via Gibson Assembly.

pCHA-FcSup-FAP0.5b.13-5E-TAA, which encodes for FAP-Fc-5E, was constructed by first modifying the portion of pCHA-FcSup-TAG that encodes the C-terminus of the construct to contain 5 additional glutamic acid codons with a modified TAA codon for termination (termed pCHA-FcSup-5E-TAA). pCHA-FcSup-5E-TAA was constructed by PCR amplifying two fragments from pCHA-FcSup-TAG. One fragment was generated by using primers pCHA-XmaI-FP and 5E-scFvFc17-R1, and the other was generated by using primers 5E-scFvFc17-F2 and 5E-scFvFc17-R2. These two segments were recombined with the pCHA-FcSup-TAG vector digested with XhoI and XmaI via Gibson Assembly. Once we obtained pCHA-FcSup-5E-TAA, we PCR amplified the FAP0.5b.13 scFv from pCHA-FcSup-FAP0.5b.13-TAG using primers FAP0.5b.13-insert-FP and FAP0.5b.13-insert-RP. We then used Gibson Assembly to recombine the scFv fragment with pCHA-FcSup-5E-TAA digested with NheI and XmaI.

pCHA-FcSup-scFv17-15E-TAA and pCHA-FcSup-FAP0.5b.13-15E-TAA (which encode for scFv-Fc17-15E and FAP-Fc-15E, respectively) were generated by first modifying the portion of pCHA-FcSup-TAG that encodes the C-terminus of the construct to contain 15 additional glutamic acid codons with a modified TAA codon for termination (termed pCHA-FcSup-15E-TAA). pCHA-FcSup-15E-TAA was generated by first PCR amplifying a gene fragment (purchased from IDT; SI Table 3) with primers 15E-insertion-new-FP and 15E-insertion-new-RP. This fragment was recombined with pCHA-FcSup-TAG digested at BglII and BamHI restriction sites via Gibson Assembly. To generate pCHA-FcSup-scFv17-15E-TAA we PCR amplified the DNA sequence for scFv17 mentioned above using primers scFv17-FP and scFv17-RP and performed Gibson Assembly to combine the amplified fragment with pCHA-FcSup-15E-TAA digested with NheI and XmaI. To generate pCHA-FcSup-FAP0.5b.13-15E-TAA, we PCR amplified the FAP0.5b.13 scFv from pCHA-FcSup-FAP0.5b.13-TAG using primers FAP0.5b.13-insert-FP and FAP0.5b.13-insert-RP and performed Gibson Assembly to recombine it with pCHA-FcSup-15E-TAA digested with NheI and XmaI.

Yeast Culture and Media
The *Saccharomyces cerevisiae* strain RJY100 used in these experiments was constructed as described previously[1]. Frozen-EZ Yeast Transformation II Kits (Zymo Research) were used to prepare and transform competent yeast, and SD-CAA and YPG media used were prepared as previously described[1, 3]. Penicillin-streptomycin 100× (penicillin: 10,000 IU and streptomycin: 10,000 µg/ml; Corning) was used during propagation and induction of yeast cultures at a final concentration of 1× in culture media.

Protein Secretion and Purification
scFv-Fcs were secreted according to previously reported methods[1, 4]. Briefly, plasmids encoding scFv-Fcs (pCHA plasmids contain a tryptophan auxotrophic marker) were transformed into Zymo competent *Saccharomyces cerevisiae* strain RJY100[1] and plated on SD-CAA solid media. Individual colonies were allowed to grow at 30 °C for 3 days. Colonies were inoculated and grown in SD-CAA liquid media supplemented with penicillin-streptomycin (pen-strep). Saturated cultures were induced in 1 L YPG supplemented with pen-strep at 20 °C for 4 days with shaking at 250–300 rpm. Following induction, cultures were pelleted by centrifuging at 3,214 rcf for 30 minutes, and cleared supernatant was passed through a 0.2 µm filter. The pH of the supernatant was neutralized by adding pH 7.4 10× PBS to a final concentration of 1×. The filtrate was passed through a column containing protein A resin (GenScript) twice, and the resin was washed three times with 1× PBS at pH 7. Protein was eluted from the resin with 7 mL 100 mM glycine at pH 3 and immediately neutralized with 0.7 mL 1 M Tris-HCl at pH 8.5. The resin was washed again with 25 mL 1× PBS at pH 7, the filtrate was passed over the column once more, and the washing and elution process was repeated. The eluant was buffer exchanged into 1× PBS pH 7.4 and concentrated using Amicon Ultra 30 kDa molecular weight cutoff devices (Millipore Sigma). Protein concentration was measured using absorbance measurements at 280 nm using a Nanodrop One device. Protein purity was assessed using 4–12% Bis Tris sodium dodecyl sulfate polyacrylamide gel electrophoresis (SDS-PAGE) gels (Invitrogen; SI Figure 1). Proteins were stored at 4 °C for up to one week, and otherwise stored by mixing protein 1:1 (vol:vol) with 100% glycerol, flash freezing in liquid nitrogen, and storing at –80 °C. Frozen protein stocks were stored in single use aliquots. To retrieve proteins from glycerol stocks, proteins were thawed at room temperature and buffer exchanged into 1× PBS pH 7.4.

FITC Conjugation
Fluorescein isothiocyanate isomer I (FITC) was obtained from Molecular Probes for fluorescently labelling proteins. FITC was dissolved in 0.1 M sodium bicarbonate to obtain a

concentration of 1–2 mg/mL just before use. ScFv-Fcs stored in 1× PBS (0.5–1 mg) were combined with FITC solution to achieve a 20:1 FITC to protein molar ratio for reaction. Reaction volume was adjusted to 250 µL with 1× PBS and 1 M sodium bicarbonate to achieve 0.1 M sodium bicarbonate final concentration for preferred reaction conditions. FITC conjugation reaction proceeded for 2 hours at 20 °C with 250–300 rpm shaking. Once the reaction finished, unbound FITC was removed first via buffer exchange using 1× PBS pH 7.4. The protein concentration and FITC to protein ratio (F/P) was monitored between centrifugation steps, and once the F/P changed by small increments after spins, labelled protein was passed through two desalting columns (7 kDa MWCO; Zeba) to ensure full removal of unbound FITC.

Mammalian Cell Culture

HepG2 cells were obtained from ATCC and cultured in low glucose DMEM (Gibco) and supplemented with 10% vol/vol fetal bovine serum (FBS; Seradigm). 100× penicillin-streptomycin (Corning) was added to a 1× final concentration in media. Luciferase reporter cell lines were constructed as described previously[5], and were obtained as a gift from the Frank Lab at the Dana Farber Cancer Institute. STAT3-, STAT1, and NFκB-luciferase cells were cultured in high glucose DMEM (Gibco) and supplemented with 10% vol/vol FBS (Seradigm). 100× penicillin-streptomycin (Corning) was added to a 1× final concentration to media. All cells were subcultured using 0.25% trypsin-EDTA (Gibco) and were maintained at 37 °C under 5% $CO_2$ without shaking.

Cytokine Stimulation

STAT3-luciferase cells were activated using human oncostatin M (OSM), STAT1-luciferase cells were activated using human interferon-gamma (IFNγ), and NFκB-luciferase cells were activated using murine tumor necrosis factor-alpha (TNFα), where each cytokine was obtained from PeproTech. HepG2 cells were stimulated with human interleukin-6 (IL-6) obtained from Gibco. For luciferase assays, luciferase reporter cells were stimulated for 5 hours at a final concentration of 10 ng/mL in media of the respective cytokine. For RT-qPCR assays, HepG2 cells were stimulated for 90 minutes at a final concentration in media of 50 ng/mL IL-6. For Western blotting experiments as in SI Figure 2, cells were stimulated for 15 minutes, where STAT3-luciferase cells were treated with 20 ng/mL OSM and HepG2 cells were treated with 30 ng/mL IL-6 in media.

Western Blotting Cell Lysates

For experiments shown in SI Figure 2, HepG2 cells and STAT3-luciferase cells were seeded in T25 flasks for overnight attachment. Each cell line was seeded in two flasks each, one for cytokine stimulation and one unstimulated for a negative control. Following cell attachment overnight, one flask of each cell line was stimulated under conditions described in "Cytokine Stimulation" section (stimulation occurred for 15 minutes where STAT3-luciferase cells were treated with 20 ng/mL OSM and HepG2 cells were treated with 30 ng/mL IL-6). Following stimulation, all cells were collected from each flask using trypsin-EDTA, and cells were washed 3 times with cold 1× PBS pH 7.4 and counted. Cells were resuspended in lysis buffer (25 mM Tris-HCl (pH 7.4), 150 mM NaCl, 1 mM EDTA, 5% glycerol, 1% NP-40, supplemented with Protease and Phosphatase inhibitor Mini Tablets, EDTA-Free (Pierce)) and lysed by sonicating for 15 seconds and incubating for 30 minutes at 4 °C on a rotary wheel. Cell lysate was cleared by centrifuging at 20,000 rcf for 20 minutes at 4 °C, and lysate was separated from cell debris and stored on ice for immediate use. Total protein concentration was measured using BCA Assay (Pierce) according to the manufacturer's instructions. 5 µg protein was loaded into SDS-PAGE gels and transferred on nitrocellulose membranes using an iBlot2 Dry Blotting System (Life Technologies). Membranes were blocked for 1 hour at room temperature in 1× TBS supplemented with 0.1% Tween-20 and 5% w/v BSA on an orbital shaker. Membranes were separately probed with antibodies for GAPDH (1:1000; Cell Signaling Technology), STAT3 (1:1,000; Cell Signaling Technology), pYSTAT3 (1:1,000; Cell Signaling Technology), and scFv-Fcs (20 µg/mL). Membranes with primary antibodies for GAPDH, whole STAT3, and pYSTAT3 used goat anti-rabbit AlexaFluor 488 (Invitrogen) as a secondary antibody, while blots with scFv-Fc primary antibodies used goat anti-human IgG Fc DyLight488 as a secondary antibody. All membranes were observed using an Azure c400 gel imager.

Lipid Synthesis and Nanoparticle Fabrication

Lipids 87-N18B[6], 113-O16B[6], EC16-80[7], 306-O16B3[8], and PBA-Q76-O16B[9] were synthesized using our previously described procedures. NanoAssemblr® (Precision NanoSystems, Inc) was used to fabricate the lipid nanoparticles. Typically, the ethanol phase was prepared by dissolving synthetic lipids (1 mg) into pure ethanol (200 uL), and the aqueous phase is either sodium acetate buffer (25 mM, pH 5.2; for 87-N18B, 113-O16B, EC16-80, and 306-O16B) or distilled water (for PBA-Q76-O16B). The two phases were then mixed by a NanoAssemblr® using the NA Ingnite NxGen cartridges to fabricate lipid nanoparticles. The ethanol was removed through dialysis against DI water (Thermo Scientific Slide-A-Lyzer™ MINI Dialysis Device, 3.5K MWCO) for 8-12 h at room temperature. Distilled water was added to the purified lipid nanoparticle aqueous solution to make the final lipid concentration 1 g/L, and the stock solution was stored at 4 °C before use. Hydrodynamic size, polydispersity index, and zeta-potential of lipid nanoparticles were measured by a Zeta-PALS particle size analyzer (Brookhaven Instruments).

Intracellular Delivery of scFv-Fcs

Lipid nanoparticles and scFv-Fcs were complexed according to experimental details summarized in SI Table 3. Volumes of both lipid (1 g/L) and scFv-Fcs (variable concentration; 0.1–10 mg/mL) were combined with enough 1× PBS (pH 7.4) to achieve the desired volume to add to wells. Lipid and scFv-Fcs combined in PBS were allowed to self-assemble for 15 minutes stationary at room temperature. After 15 minutes, the desired volume was added to media in each well, carefully as to not disturb the cells.

Flow Cytometry Analyses

HepG2 cells (20,000 cells per well), STAT3-luciferase cells (10,000 cells per well), and STAT1-luciferase cells (10,000 cells per well) were seeded in 48-well plates 24 hours prior to delivery. Additional wells were seeded for cell-only and protein-only controls for flow cytometry gating analysis. Nanocomplexes were assembled as described above, where FITC-labelled scFv-Fc and lipid masses were used depending on the type of experiment (SI Table 3). Protein-only controls were prepared by diluting the mass of FITC-scFv-Fcs used in the experiment in 1× PBS without lipid. 24 hours after delivery, cells were washed three times with 1× PBS containing 20,000 U/mL heparin to remove particles bound to the cell surface[10]. The cells were then incubated in 0.05% phenol-free trypsin-EDTA (Gibco) diluted with 1× PBS (diluted 7:3, trypsin to PBS) for 10 minutes at 37 °C. Cell suspensions were aliquoted either to microcentrifuge tubes or V-bottom 96-well plates for flow cytometry analysis using an Attune NxT Flow Cytometer (Life Technologies). Unless otherwise stated, 10,000 events were collected for each sample on the flow cytometer. Samples were analyzed using FlowJo software, where viable cells were gated with cell-only control samples, and baseline for FITC-positive cells were set with protein-only control samples.

Size and Encapsulation Efficiency Characterizations

scFv-Fcs were complexed with PBA-Q76-O16B in a 1:8 protein to lipid ratio in 1× PBS as described above and in SI Table 3 in order to measure the particle size with dynamic light scattering (DLS; ZataPALS, Brookhaven Instruments). Encapsulation efficiency was determined by separating the nanocomplexes from the supernatant by centrifugation at 21,000 rcf for 45 minutes at 4 °C. Protein concentration of supernatant was assessed with the BCA Assay (Pierce), and the percent encapsulated protein was determined by mass balance.

Cell Viability Assays

Cell viability was assessed by first seeding either 3,000 cells per well of luciferase reporter cell lines or 6,000 cells per well of HepG2 cells in white-walled 96-well flat bottom plates. 24 hours after cell seeding, nanocomplexes were assembled as described above and in SI Table 3 and added to necessary wells. Lipid only wells used PBA-Q76-O16B diluted in 1× PBS. Control wells were left untreated with nanoparticles for viability calculations. 24 hours after delivery, the CellTiter-Fluor Cell Viability Assay (Promega) was used according to the manufacturer's instructions. Fluorescent data was recorded on a SpectraMax i3x Microplate Reader (Molecular Devices).

Luciferase Assays

24 hours prior to delivery, luciferase reporter cells were seeded at a density of 3,000 cells per well in white-walled 96-well flat bottom plates. Nanocomplexes were assembled as described above and in SI Table 3 such that each well would receive 10 nM scFv-Fc final concentration per well complexed with PBA-Q76-O16B in a 1:8 protein to lipid mass ratio. 24 hours after delivery, wells with delivered nanoparticles, as well as a set of wells without nanoparticles for controls, were stimulated with 10 ng/mL of the respective cytokine, depending on which cell line was being used (see above for compatible cytokines). After 5 hours of stimulation, BrightGlo Luciferase Assay System (Promega) was used according to the manufacturer's instructions to measure luciferase expression. CellTiter-Fluor Cell Viability Assay (Promega) was used to measure cell viability in each well according to the manufacturer's instructions for multiplexing with luciferase detection assays. Fluorescent and luminescent data was recorded on a SpectraMax i3x Microplate Reader. Luminescent data was normalized to viable cell number from fluorescent data and further normalized relative to stimulated controls.

RT-qPCR Assays

24 hours prior to delivery, HepG2 cells were seeded at a density of 20,000 cells per well in 48-well plates. Nanocomplexes were assembled as described above and in SI Table 3 such that each well would receive 10 nM scFv-Fc final concentration per well complexed with PBA-Q76-O16B in a 1:8 protein to lipid mass ratio. Some wells of cells did not have nanoparticles delivered and served as controls for analysis. 24 hours after delivery, all wells containing nanoparticles, in addition to some control wells without nanoparticles, were washed once with 1× PBS and incubated with media containing 50 ng/mL IL-6 for 90 minutes. Following cytokine stimulation, total RNA was extracted from each well of cells using TriZol (Life Technologies) according to the manufacturer's instructions. 500 ng RNA was then reverse transcribed using SuperScript IV (SSIV) First-Strand Synthesis System (Invitrogen). cDNA produced was used with PowerUp SYBR Green Mater Mix (Applied Biosystems) and primers listed in SI Table 5 for quantitative PCR, where data was collected on a QuantStudio5 Real-Time Thermocycler (Applied Biosystems). TriZol, SSIV First-Strand Synthesis System, and PowerUp SYBR Green Mater Mix were all used according to the manufacturer's instructions. Raw data was analyzed using QuantStudio Design and Analysis Software and further analyzed using the $\Delta\Delta C_T$ method[11, 12], where HPRT was used as the housekeeping gene for all calculations.

### Supplementary Figures

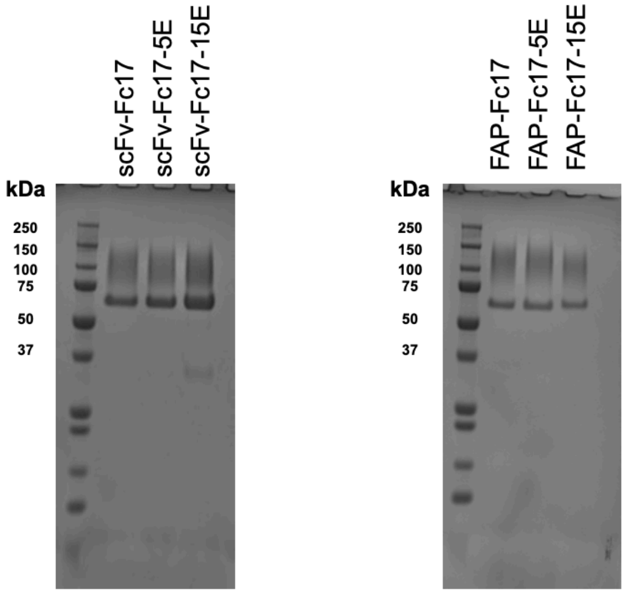


**SI Figure 1.** SDS-PAGE gels depicting purified proteins used for this study. Smear pattern is a result of glycosylation from secretory pathway in yeast cells, and lower band is monomeric scFv-Fc[1].

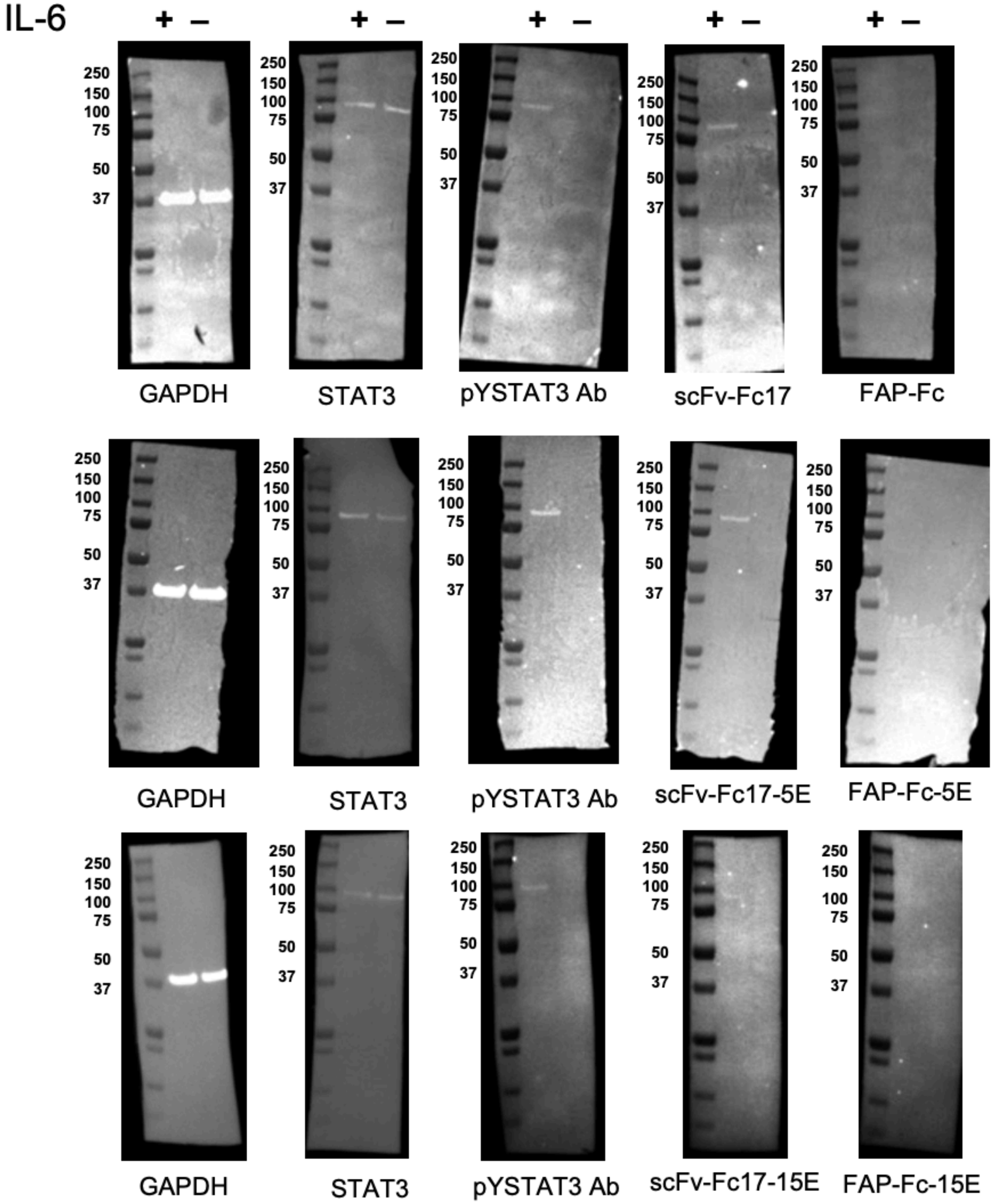


**SI Figure 2.** Immunoblots showing specific binding of scFv-Fc17 constructs to pYSTAT3 in HepG2 cells. HepG2 cells were seeded in T25 flasks overnight to attach. The following morning, cells were stimulated with 30 ng/mL IL-6 for 15 minutes. Cells were lysed and 5 µg lysate was loaded into SDS-PAGE gels and blotted for GAPDH, whole STAT3, pYSTAT3 using a commercial antibody (Ab), scFv-Fc17 constructs (20 µg/mL), and FAP-Fc constructs (20 µg/mL).

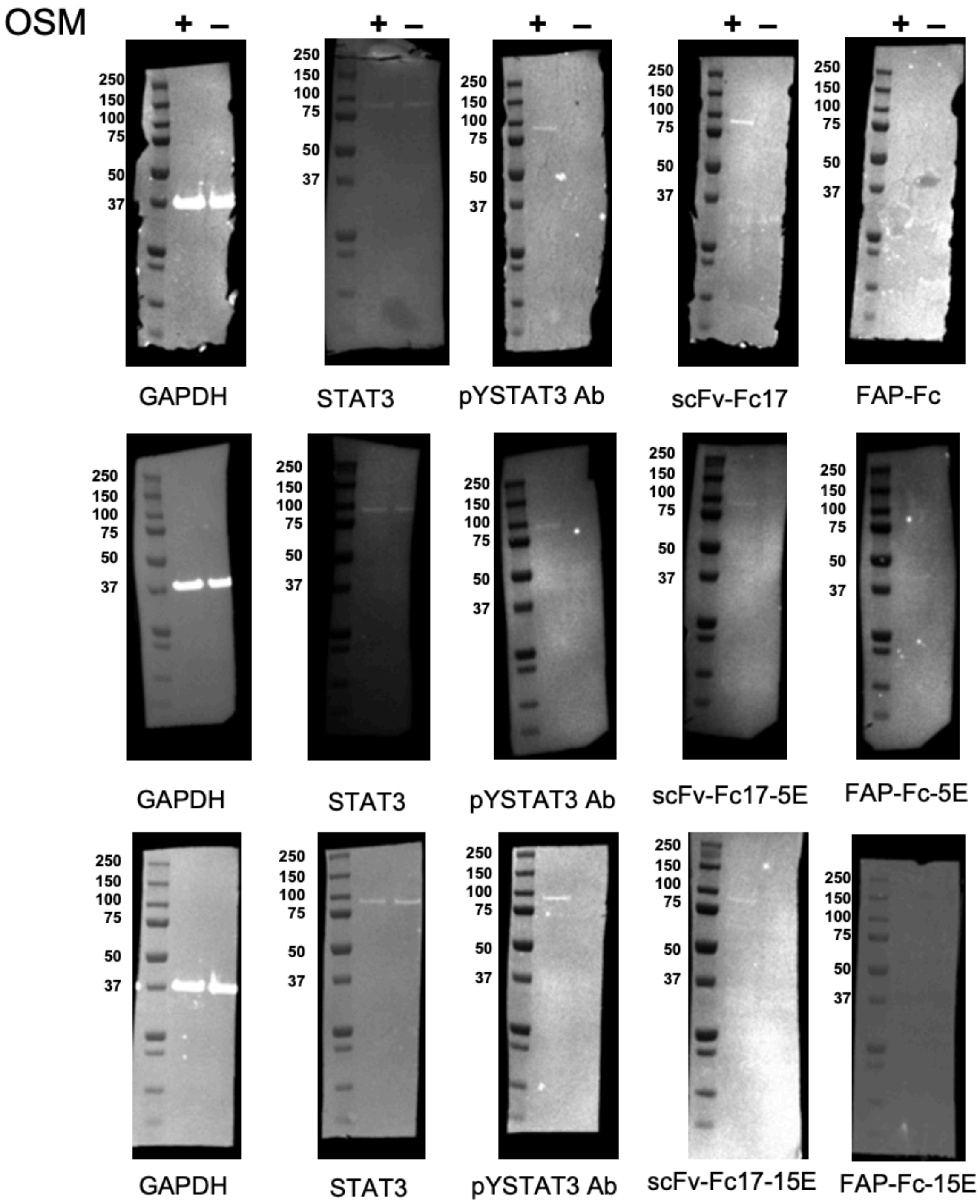


**SI Figure 3.** Immunoblots showing specific binding of scFv-Fc17 constructs to pYSTAT3 in STAT3-luciferase cells. STAT3-luciferase cells were seeded in T25 flasks overnight to attach. The following morning, cells were stimulated with 20 ng/mL OSM for 15 minutes. Cells were lysed and 5 µg lysate was loaded into SDS-PAGE gels and blotted for GAPDH, whole STAT3, pYSTAT3 using a commercial antibody (Ab), scFv-Fc17 constructs (20 µg/mL), and FAP-Fc constructs (20 µg/mL).

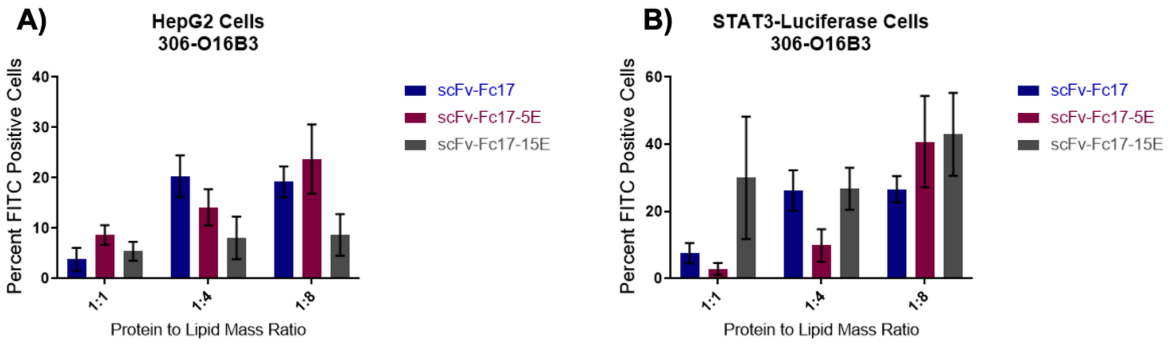


**SI Figure 4.** Flow cytometry analysis of changes in protein to lipid mass ratio for delivery. A) HepG2 cells and B) STAT3-luciferase cells were incubated with 10 nM scFv-Fcs complexed at indicated mass ratios with lipid 306-O16B3 for 24 hours. Following incubation, cells were washed three times with 1× PBS containing 20 U/mL heparin and then analyzed via flow cytometry for detection of FITC-labelled scFv-Fcs. Data shown is a mean three technical replicate wells where error is the standard deviation from the mean (n = 3).

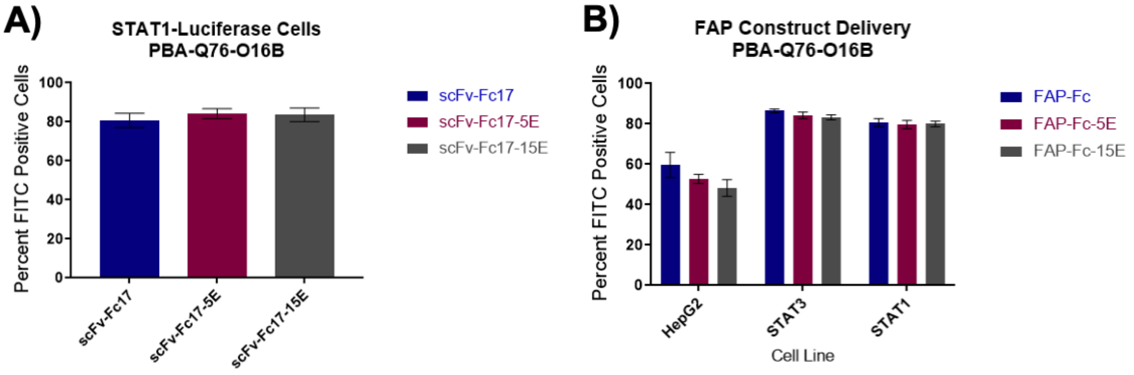


**SI Figure 5.** Control cell lines and proteins. A) STAT1-luciferase cells were incubated with indicated 10 nM scFv-Fcs complexed in a 1:8 protein to lipid mass ratio with PBA-Q76-O16B for 24 hours. Following incubation, cells were washed three times with 1× PBS containing 20 U/mL heparin and then analyzed via flow cytometry for detection of FITC-labelled scFv-Fcs. Data shown is a mean of two independent experiments each containing three technical replicate wells where error is the standard deviation from the mean (n = 6). B) Specified cell lines were incubated with indicated scFv-Fcs complexed in a 1:8 protein to lipid mass ratio with PBA-Q76-O16B for 24 hours. Following incubation, cells were washed three times with 1× PBS containing 20 U/mL heparin and then analyzed via flow cytometry for detection of FITC-labelled scFv-Fcs. Data shown is a mean of two independent experiments each containing three technical replicate wells where error is the standard deviation from the mean (n = 6).

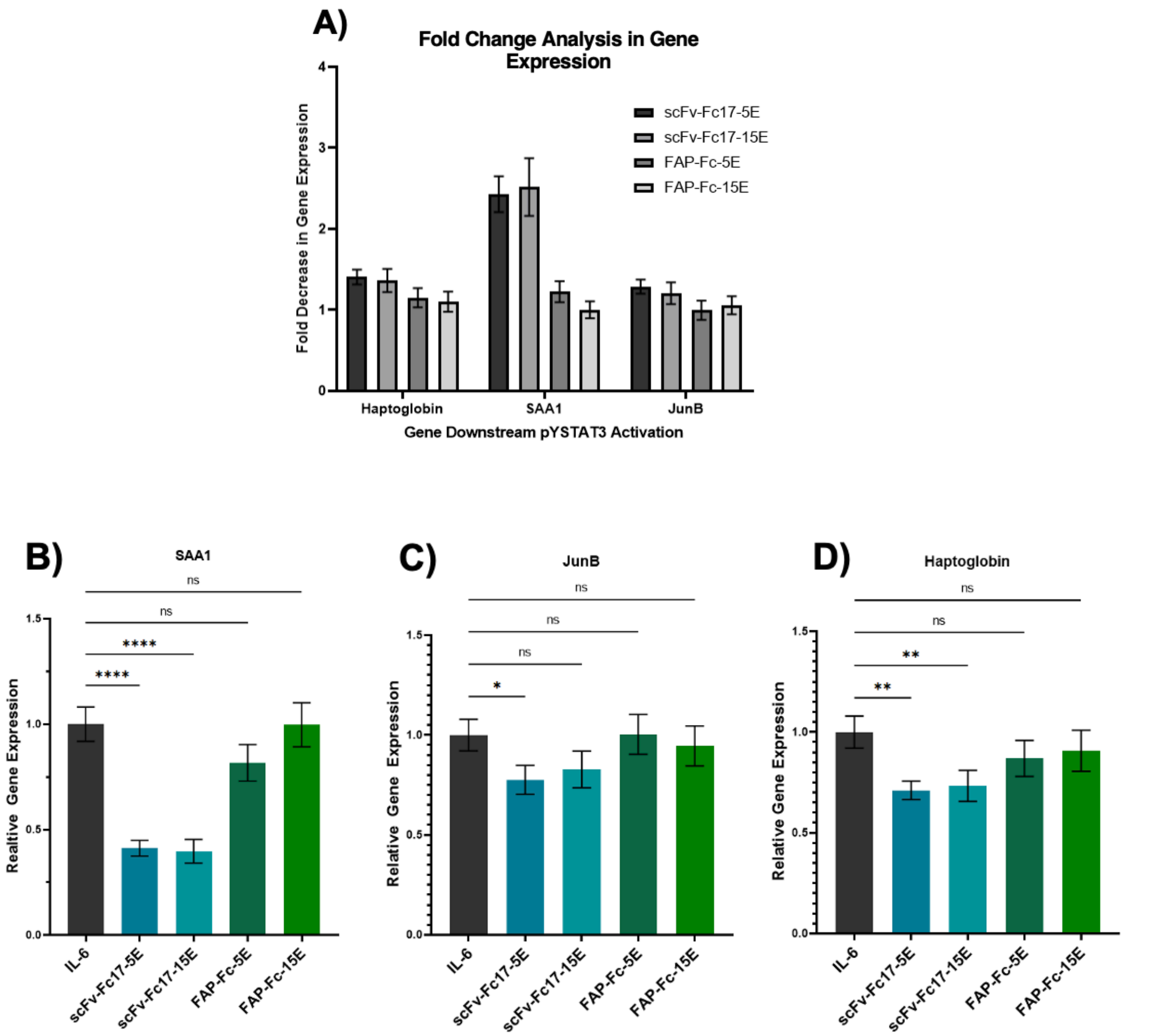


**SI Figure 6.** Alternative representation of data shown in main text Figure 6. A) Data shown represents the fold decrease in gene expression from the IL-6 stimulated control (where cells were treated only with IL-6 and no nanoparticles). B–D) Data shown is the gene expression normalized to the IL-6 only control, and error is represented by the standard deviation propagated through analysis (n = 3). Statistical analysis is from a one-way ANOVA where **** shows $p \leq 0.0001$, ** shows $p \leq 0.01$, * shows $p \leq 0.05$, and ns shows no significance.

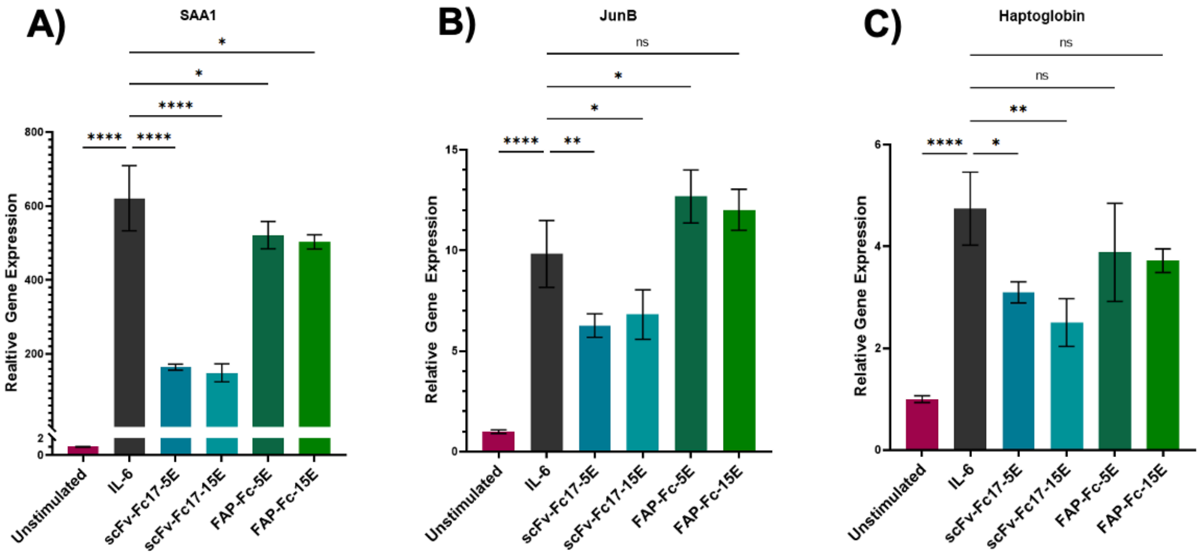


**SI Figure 7.** Repeat RT-qPCR experimental data. Analysis for A) SAA1, B) JunB, and C) haptoglobin. HepG2 cells were incubated with indicated scFv-Fcs at a 10 nM final concentration complexed with lipid PBA-Q76-O16B in a 1:8 mass ratio for 24 hours in media. Cells were stimulated with 50 ng/mL IL-6 for 90 minutes to activate pYSTAT3 pathway. Total RNA was extracted with TriZol, reverse transcribed with SSIV First Strand Synthesis System, and amplified using PowerUp SYBR Green Master Mix using primers listed in *Materials and Methods*. qPCR data was analyzed using the $\Delta\Delta C_T$ method using HPRT as a housekeeping gene. Data shown is the relative gene expression normalized to the unstimulated control, and error is represented by the standard deviation propagated through analysis (n = 3). Statistical analysis is from a one-way ANOVA where **** shows $p \leq 0.0001$, ** shows $p \leq 0.01$, * shows $p \leq 0.05$, and ns shows no significance.

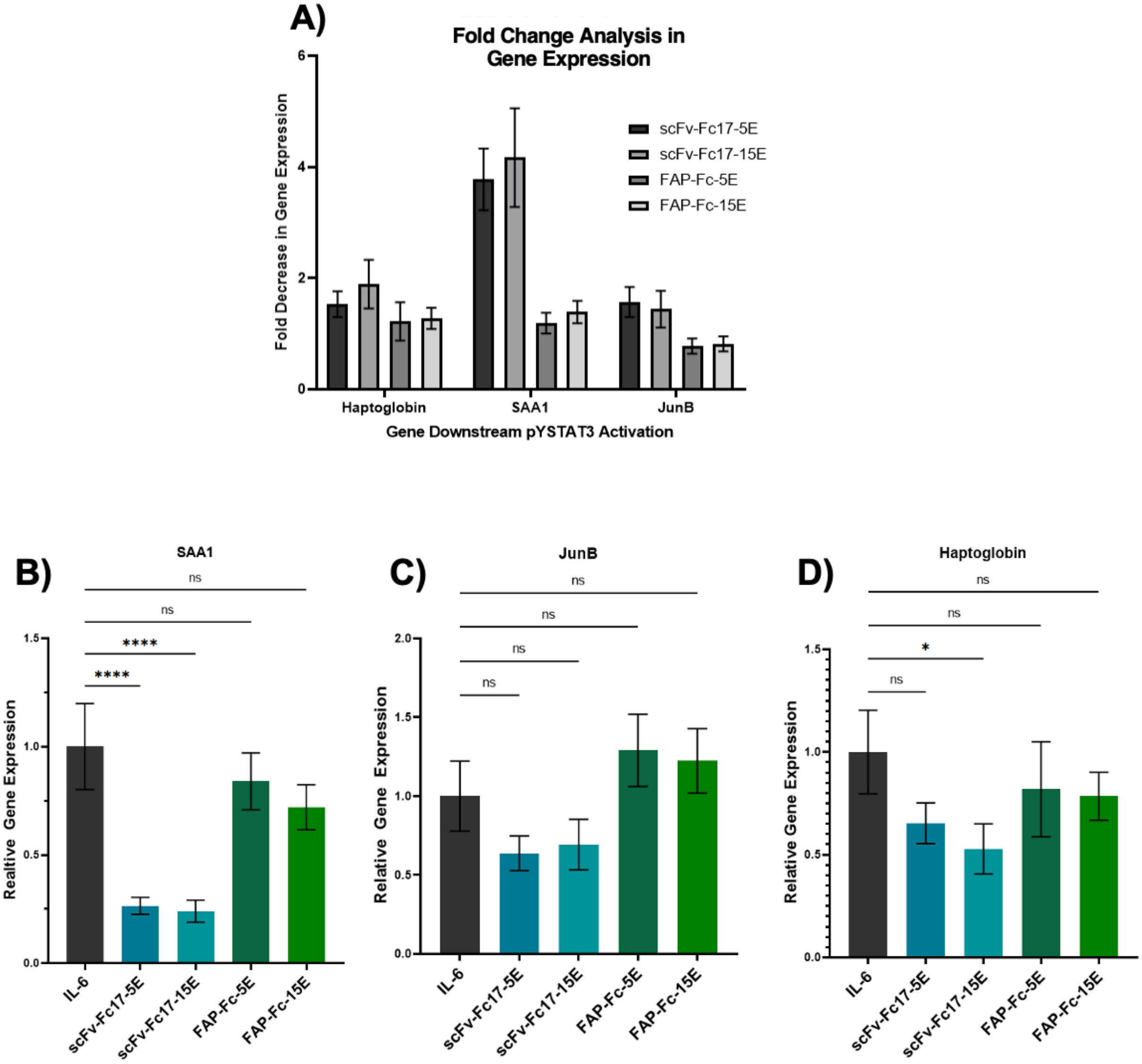


**SI Figure 8.** Alternative representation of data shown in SI Figure 7. A) Data shown represents the fold decrease in gene expression from the IL-6 only stimulated control where cells were treated only with IL-6 and no nanoparticles. B–D) Data shown is the gene expression normalized to the IL-6 only control, and error is represented by the standard deviation propagated through analysis (n = 3). Statistical analysis is from a one-way ANOVA where **** shows $p \leq 0.0001$, ** shows $p \leq 0.01$, * shows $p \leq 0.05$, and ns shows no significance.

### Supplementary Tables

**SI Table 1.** scFv sequences.

| | |
|---|---|
| scFv17 | GAACTGGTGTTGACCCAAACACCGAGTCCTGTCAGTGCGGTCG<br>TCGGGGGGACTGTGACTATCAACTGTCAAAGTTCACAGAGCGT<br>GTGGGGAAATAATAGGTTAAGCTGGTACCAGCAGAAGCCAGGT<br>CAACCACCACGTTTACTTATGTACTATGCTTCTAACCTTGCCTCC<br>GGAGTCTCATCCAGATTTAAAGGGAGTGGATCTGGCACGCAGT<br>TTACTTTAACCATAAGCGATGTACAGTGCGATGACGCAGCAACA<br>TACTACTGCCAAGGCGGCTTCGAGTGCAGCGGCGGAGATTGC<br>GTCGGTTTTGGAGGGGGGACTGAGTTAGAAATCTTGGGCGGGA<br>GCTCTAGGTCTTCCTCCAGTGGAGGCGGAGGTTCAGGAGGCG<br>GCGGCCAGTCCGTCGAAGAATCCGGTGGTAGGCTAGTCGCTC<br>CCGGAGGAAGTCTGACCCTTACATGTACCGTGTCCGGGATCGA<br>CCTGTCATCCGACGCAATGAGTTGGGTCAGGCAGGCGCCTGGT<br>AAAGGATTGGAATGGATCGGAACGATATATGGCAGCGCAGGTA<br>CTTACTACGCAACTTGGGCTAAGGGTCGTTTCACTATCAGTAAA<br>ACGTCAACGACTGTTGACTTAAAGATGACTTCTTTAACGACGGA<br>AGACACCGCCACATACTTCTGCACACGTGCCTTCTCCAATACTA<br>GACTAGATCTATGGGGCCAGGGGACCTTGGTTACCATTTCCTC<br>TGGTGCCGGAGGGGCGTCTGGAGGAGGAGGTTCCGGTGGTG<br>GTGGCTCA |
| FAP0.5b.13 | GATATCCAGATGACCCAGTCCCCGAGCTCCCTGTCCGCCTCTG<br>TGGGCGATAGGGTCACCATCACCTGCCGTGCCAGTCAGTACTC<br>TGGTTCTGTAGCCTGGTATCAACAGAAACCAGGAAAAGCTCCG<br>AAGCTTCTGATTTACTCTGCATCCTACCTCTACTCTGGAGTCCC<br>TTCTCGCTTCTCTGGTAGCCGTTCCGGGACGGATTTCACTCTGA<br>CCATCAGCAGTCTGCAGCCGGAAGACTTCGCAACTTATTACTGT<br>CAGCAAGGTTCTGCTCTGATCACGTTCGGACAGGGTACCAAGG<br>TGGAGATCAAAGGTACTACTGCCGCTAGTGGTAGTAGTGGTGG<br>CAGTAGCAGTGGTGCCGAGGTTCAGCTGGTGGAGTCTGGCGG<br>TGGCCTGGTGCAGCCAGGGGGCTCACTCCGTTTGTCCTGTGCA<br>GCTTCTGGCTTCAACCTCGGTGGTTACTACATGCACTGGGTGC<br>GTCAGGCCCCGGGTAAGGGCCTGGAATGGGTTGCATCTATTTA<br>CCCTGGTTCTGGCGGTACTTACTATGCCGATAGCGTCAAGGGC<br>CGTTTCACTATAAGCGCAGACACATCCAAAAACACAGCCTACCT<br>ACAAATGAACAGCTTAAGAGCTGAGGACACTGCCGTCTATTATT<br>GTGCTCGCGGTTGGTCTTCTGCTATGGACTACTGGGGTCAAGG<br>AACCCTGGTCACCGTCTCCTCGGGACCCGGG |

**SI Table 2.** Primer sequences for cloning scFv-Fcs.

| | |
|---|---|
| scFv17-FP | TACCCATACGACGTTCCAGACTACGCTGCTAGCGAACTGGTGT TGACCCAAACACCGA |
| scFv17-RP | TTTGTCGGAACTTTTAGGTTCTACCCCGGGTGAGCCACCACCA CCGGAACCTCCTC |
| 5E-scFvFc17-F2 | GGTAAGAGATCTGAAGAAGAGGAAGAGTAAGGTGGAGGAGGC TCTGGTGGAGG |
| 5E-scFvFc17-R1 | CCTCCACCAGAGCCTCCTCCACCTTACTCTTCCTCTTCTTCAGA TCTCTTACC |
| 5E-scFvFc17-R2 | ACATCTACACTGTTGTTATCAGATTTCGCTCGAG |
| pCHA-XmaI-FP | ACGCTGCTAGCCAGTCGCATATGACTGAGCCCGGG |
| FAP0.5b.13-insert-FP | CTTACCCATACGACGTTCCAGACTACGCTGCTAGCGATATCCA GATGACCCAGTC |
| FAP0.5b.13-insert-RP | TTTTGTCGGAACTTTTAGGTTCTACCCCGGGTCCCGAGGAGAC GGTGACCAGG |
| 15E-insertion-new-FP | TATGCACGAAGCCTTACACAATCACTACACACAAAAGTCATTAT CCTTGTCTCCAG |
| 15E-insertion-new-RP | TAGTTGTCAGTTCCTGCAAGTCTTCTTCGGAGATAAGCTTTTGT TCTGCACGCGTG |

**SI Table 3.** 15E insert sequence.

| | |
|---|---|
| 15E gene fragment | ACAAAAGTCATTATCCTTGTCTCCAGGTAAGAGATCTGAGGAGG AAGAGGAAGAGGAGGAGGAGGAGGAAGAAGAGGAAGAGTAAG GTGGAGGAGGCTCTGGTGGAGGCGGTAGCGGAGGCGGAGGG TCGGGATCCACGCGTGCAGAACAAAAGCTTATC |

**SI Table 4.** scFv-Fc and lipid masses used in different experiments.

| Protein to Lipid Mass Ratio | Lipid Mass (µg) | Protein Mass (µg) | Final Protein Concentration in Media (nM) | Volume added to wells (µL) |
|---|---|---|---|---|
| Flow cytometry lipid screen (48-well plate) | | | | |
| 1:1 | 1 | 1 | 35 | 50 |
| Flow cytometry protein to lipid mass ratio changes (48-well plate) | | | | |
| 1:1 | 0.25 | 0.25 | 10 | 20 |
| 1:2 | 0.50 | 0.25 | 10 | 20 |
| 1:4 | 1 | 0.25 | 10 | 20 |
| 1:8 | 2 | 0.25 | 10 | 20 |
| 2:1 | 0.125 | 0.25 | 10 | 20 |
| 4:1 | 0.0625 | 0.25 | 10 | 20 |
| 8:1 | 0.03125 | 0.25 | 10 | 20 |
| Luciferase Assays (96-well plate) | | | | |
| 1:8 | 1 | 0.125 | 10 | 10 |
| RT-qPCR experiments (48-well plate) | | | | |
| 1:8 | 2 | 0.25 | 10 | 20 |
| Cytotoxicity (96-well plate) | | | | |
| 1:8 | 1 | 0.125 | 10 | 10 |
| Size and Encapsulation Efficiency (1.7 mL tubes) | | | | |
| 1:8 | 10 | 1.25 | N/A | 100 uL (total volume in tube) |

**SI Table 5.** qPCR primer sequences.

| | |
|---|---|
| HPRT-FP | TCAGGCAGTATAATCCAAAGATGGT |
| HPRT-RP | AGTCTGGCTTATATCCAACACTTCG |
| Haptoglobin-FP | CCAGAGGCAAGACCAACCAA |
| Haptoglobin-RP | GCAGCCGTCATCTGCGATA |
| SAA1-FP | GAGGCTTTTGATGGGGCTCG |
| SAA1-RP | CTGGATATTCTCTCTGGCATCGGT |
| JunB-FP | CGGCGGTGGCGGCAGCTACTTTTC |
| JunB-RP | GGGGGTGTCACGTGGTTCATCTTG |
